## Supplemental Materials Legends for "High performance enrichment-based genome sequencing to support the investigation of hepatitis A virus outbreaks"

### **Supplemental tables and figures**

**Table S1.** Resultant pan-HAV oligos designed for capture of all six human subgenotypes of HAV.

**Table S2.** GenBank accession numbers for sequences used to design pan-HAV oligos and associated subgenotype and collection metadata. Date of sample collection, where available, is formatted as DD-MM-YYYY.

**Table S3.** GenBank accession numbers for sequences used for *in silico* prediction of the pan-HAV oligo design efficiency, including associated subgenotype and collection metadata. Date of sample collection, where available, is formatted as DD-MM-YYYY.

**Table S4.** Summary of positive and negative controls from the optimisation experiment and resultant genome coverage results. Undetermined (Undet.) Cτ values occur when the target is not present or is present below the limit of detection of the assay used. Copy number is the genome copies/µL quantified from the RNA extract as determined by the ISO 15216-1 HAV quantification real-time RT-PCR protocol. Mean RD is calculated as the average number of deduplicated reads covering one position in the genome. Coverage reported here is the percentage of genome positions with a depth of 10 or more reads.

- Indicates no reads from the corresponding index were present in the sequence data.

**Figure S1.** Read length distribution of spike-in sample libraries measured before quality control.
