## Supplementary figures and images for "High performance enrichment-based genome sequencing to support the investigation of hepatitis A virus outbreaks"

### Supplemental Figure 1

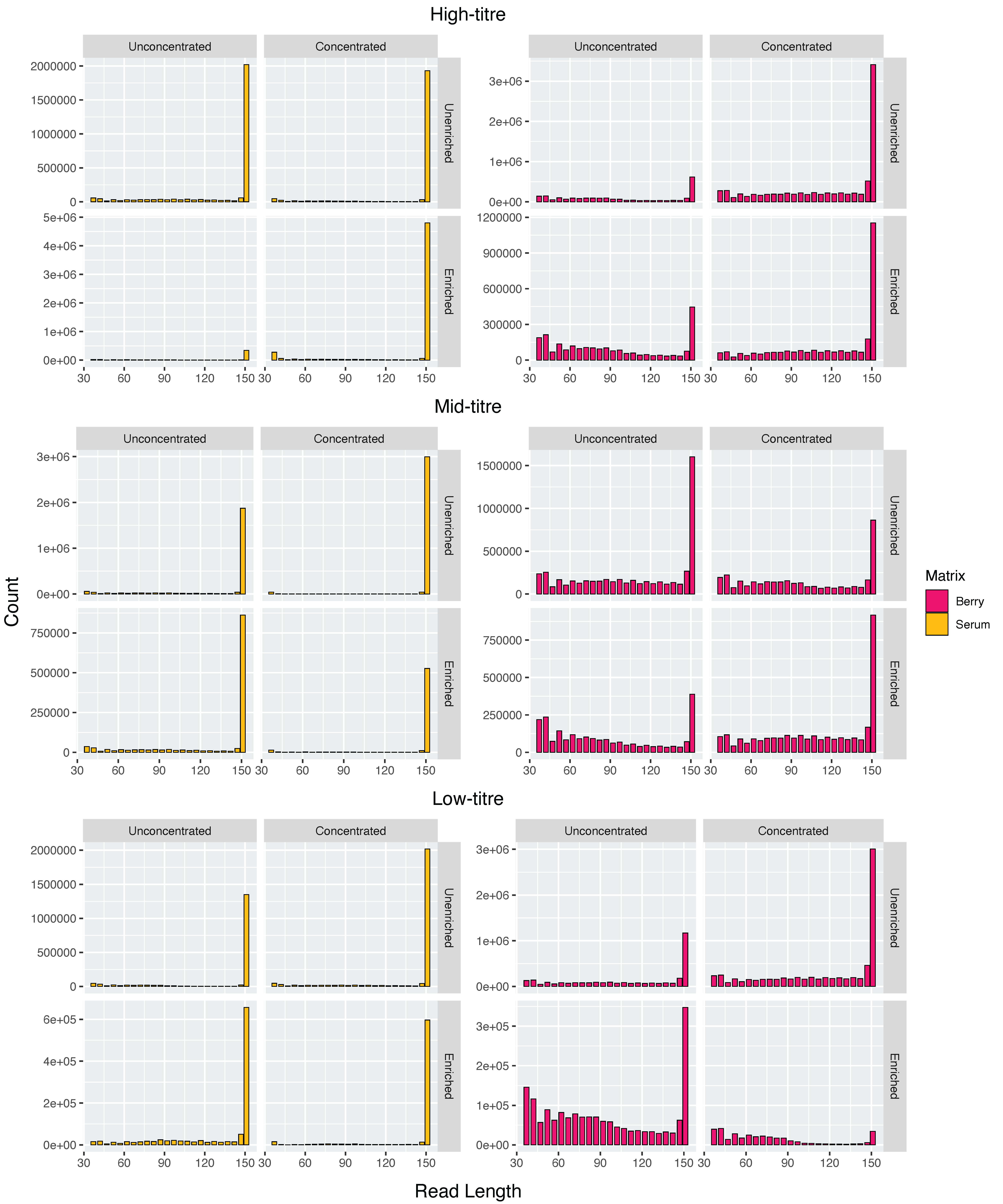
